# Age-dependent gut microbiota dynamics and their association with male fitness traits in *Drosophila melanogaster*

**DOI:** 10.1101/2025.05.22.655607

**Authors:** Zahida Sultanova, Handan Melike Dönertaş, Alejandro Hita, Prem Aguilar, Berfin Dag, José Ignacio Lucas-Lledo, Amparo Latorre, Pau Carazo

**Affiliations:** School of Biological Sciences, University of East Anglia, Norwich, UK; Leibniz Institute on Aging – Fritz Lipmann Institute (FLI), Beutenbergstrasse 11, 07745 Jena, Germany; Cluster of Excellence Balance of the Microverse, Friedrich Schiller University Jena, Jena, Germany; Cavanilles Institute of Biodiversity and Evolutionary Biology, Valencia, Spain; CIBIO Research Centre in Biodiversity and Genetic Resources, InBIO, Universidade do Porto, Porto, Portugal; BIOPOLIS Program in Genomics, Biodiversity and Land Planning, CIBIO, Campus de Vairão, 4485-661 Vairão, Portugal; Departamento de Biologia, Faculdade de Ciências da Universidade do Porto, Porto, Portugal; Laboratory of Computational Biology, VIB Center for AI & Computational Biology (VIB.AI), Leuven, Belgium; VIB-KU Leuven Center for Brain & Disease Research, Leuven, Belgium; Department of Human Genetics, KU Leuven, Leuven, Belgium; Institute for Integrative Systems Biology, Universitat de Valencia - CSIC, Valencia, Spain

## Abstract

Growing evidence suggests that the gut microbiota plays a key role in shaping life history in a wide range of species, including well-studied model organisms like *Drosophila melanogaster.* Although recent studies have explored the relationship between gut microbiota and female life history, the link between gut microbiota and male life history remains understudied. In this study, we explored the role of gut microbiota in shaping male life history traits by correlating variation in life history traits across genetically homogeneous isolines with their naturally occurring gut microbiota. Using 22 isolines from the *Drosophila melanogaster* Genetic Reference Panel (DGRP), we measured lifespan, early/late-life reproduction, and early/late-life physiological performance. We characterized the gut microbiota composition in young (5 days old) and old (26 days old) flies using 16S rDNA sequencing. We observed significant variation in male life history traits across isolines, as well as age-related changes in gut microbiota composition. Using machine learning, we showed that gut microbiota composition could predict the age of the organisms with high accuracy. Associations between gut microbiota and life history traits were notable, particularly involving the *Acetobacter* genus. In early life, the abundance of *Acetobacter ascendens* was associated with functional aging, while *Acetobacter indonesiensis* was linked to reproductive senescence. In late life, higher abundances of *A. ascendens* and *Acetobacter pasteurianus* were negatively associated with lifespan. These findings highlight the potential role of gut microbiota, especially the *Acetobacter* genus, in male fitness and aging.

## Introduction

Understanding the mechanisms of aging and reproduction is crucial to unravel the complex dynamics of life history evolution [1]. An emerging line of research highlights the role of gut microbiota [2,3], which significantly influences host lifespan, healthspan, reproduction, and behavior [4–6]. These effects can be mediated by extending metabolic capabilities beyond the host genome and modulating the immune system [7]. Notably, microbiota transfer from young to old individuals can rejuvenate hosts, enhancing both lifespan and late-life reproduction [3,8–12]. Together, these findings position gut microbiota as a central component of life history evolution [7].

The existing research on life history evolution has focused predominantly on females, resulting in a pronounced gender bias [13]. This has led to greater emphasis on the connection between gut microbiota and female life history, while male life history remains comparatively underexplored. For example, the effects of gut microbiota on female lifespan and reproduction, and to some extent male lifespan, have been well studied in *Drosophila melanogaster* [14–17] and *Caenorhabditis elegans* [18,19]. In contrast, their role in male reproduction is far less understood [20,21].

The relatively few studies exploring the influence of gut microbiota on male fitness suggest notable effects on mating behavior and reproductive success [20,22]. For example, Morimoto et al. (2017) infected *D. melanogaster* males with *Acetobacter pomorum* or *Lactobacillus plantarum*, two of the five most abundant bacterial species in wild flies [23] and known to affect *D. melanogaster* physiology and behavior [24]. Males infected with *L. plantarum* exhibited longer mating durations and caused the females to produce more offspring in the short term, while females mated with males infected with *A. pomorum* were less likely to produce viable offspring. Likewise, Ami et al. (2010) studied how gut microbiota can affect the mating behavior of the Mediterranean fruit fly (*C. capitata*). First, they damaged the gut bacterial community structure of males by sterilizing them with radiation. Then, they found that regenerating the original microbiota community of males, by feeding them with a bacteria-enriched diet, enhanced their mating performance compared to controls [22]. Finally, Heys et al. (2020) found that an intact microbiota is essential for old males to attract females in *Drosophila pseudoobscura* fruit flies [25]. Altogether, these findings show that gut microbiota is an essential factor that affects male fitness.

Our aim in this study was to explore the potential role of gut microbiota in shaping male life history traits by examining life history traits and gut microbiota across two ages of male *D. melanogaster*. First, we measured key traits in male fruit flies from 22 different DGRP inbred isolines, including life-history traits (lifespan and early/late-life reproduction) and physiological performance traits (early/late-life anti-predatory escape ability). Second, we characterized the early and late life gut microbiota of these isolines and investigated how gut microbiota composition changed with age. Finally, we explored the potential link between these male life history traits and gut microbiota composition.

## Materials & Methods

### Experimental population

As focal flies, we used flies from the *D. melanogaster* genetic reference panel (DGRP, see [26]). Hence, individuals within each isoline can be considered as clones. Using DGRP isolines (many individuals with the same genotype) instead of wild-type flies allowed us to characterize the life history traits of different genotypes in standard conditions, while characterizing the early and late life gut microbiota associated with these same genotypes. We fed the DGRP flies with Bloomington Drosophila Stock Centre Cornmeal Food (15.9 g yeast, 9.2 g soy flour, 67.1 g yellow cornmeal, 5.3 g agar, 70 mL light corn syrup and 4.4 mL propionic acid per Litre). We maintained each isoline inside a bottle with 75 mL of food until the life history assays. We also maintained sparkling poliert (*spa*) flies to be used as standard competitors (males) and mating partners (females) in the mating assays. We fed *spa* flies with a diet that contains 40 g yeast, 50 g sugar, 10 g soy flour, 60 g corn flour, 10 g agar, 3 g nipagin and 5 mL propionic acid per Litre.

### Life history assays

We set up replicate vials containing 10 males in same-sex groups for each of the 22 different DGRP isolines that were randomly chosen among the ones without *Wolbachia* infection. However, we found that isolines 492 and 427 were infected by Wolbachia **(Figure 2c)**. It is unclear whether these infections were present before arriving at our laboratory or originated during maintenance in our laboratory. We transferred these flies to new vials with fresh food once a week throughout their lifespan (or until sacrificed, see below), and checked mortality 5-6 days a week by recording the number of dead individuals in each experimental vial. Density within vials was kept constant between 8-11 individuals (by re-distributing the flies from the low-density vials among the remaining vials) until the density decreased inevitably later in the experiment as the last flies died **(Figure 1)**. To estimate the reproductive success of focal males, we measured the relative paternity of all experimental males competing against standard rivals at two different time points: early (4 days old) and late (25 days old) in life. Namely, we introduced 10 focal males with 10 *spa* males and 10 *spa* females into new vials and let them interact and lay eggs for 24 hours. At the end of this period, we recovered the focal males belonging to the isolines and discarded the individuals with the *spa* mutation. Following the first reproductive assay, on day 5, we sacrificed 15-20 males per isoline for gut dissection and kept the remaining flies for life history characterization. We incubated the eggs from the first reproductive assay and left them to develop into adults for 16 days, froze them, and then counted the proportion of wt/*spa* offspring as a measure of their reproductive success. We repeated this procedure on day 25, and again sacrificed 15-20 males per isoline for gut microbiota analyses on day 26. We calculated reproductive aging by subtracting average late life reproductive success from average early life reproductive success per isoline.

**Figure 1:**
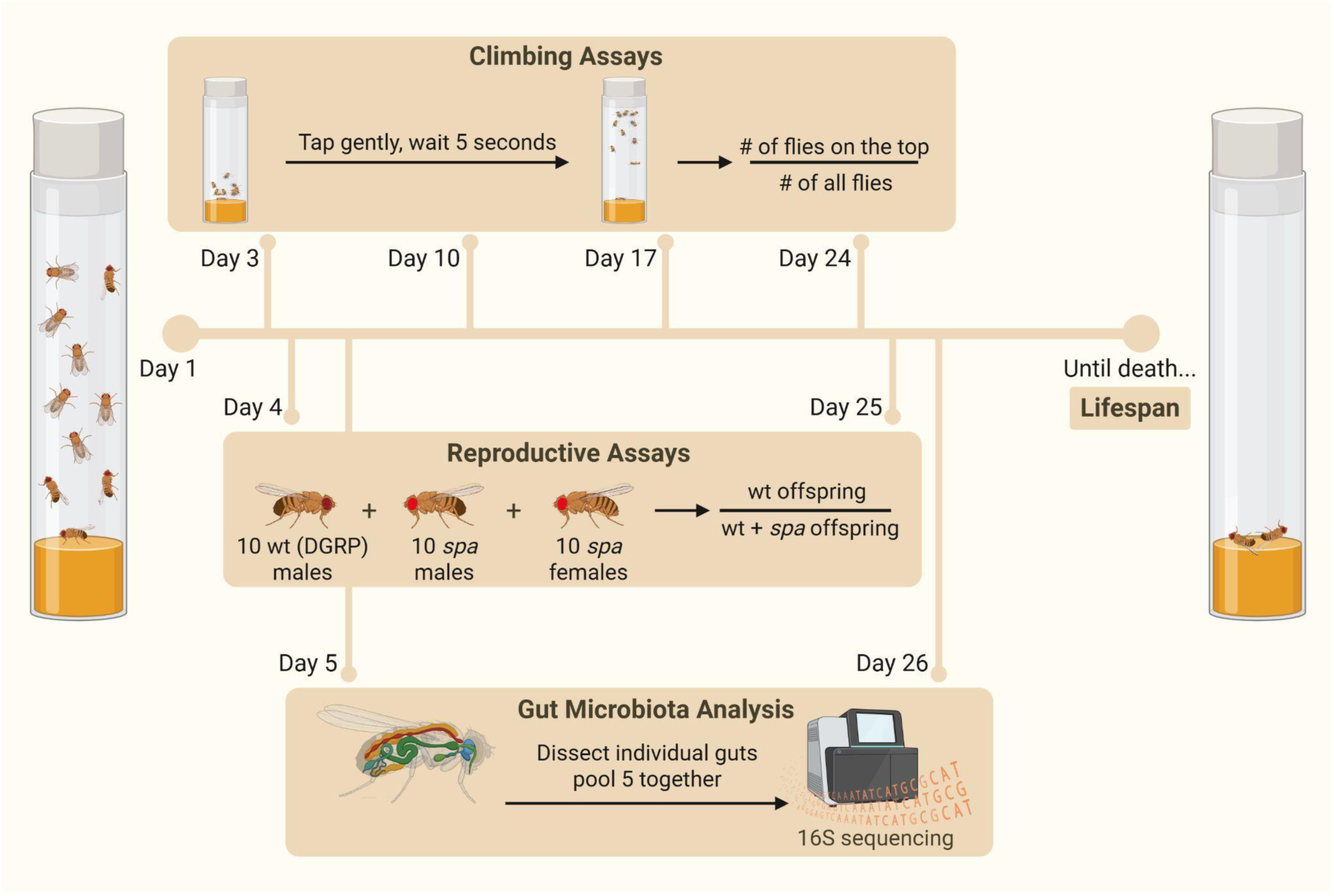
Experimental design. We monitored replicate vials that contained 10 males in same-sex groups for each of the 22 different DGRP isolines. For climbing assays, we tapped each vial and measured the proportion of flies that were able to climb to the top in 5 seconds. We measured climbing speed once a week for the first 4 weeks. For reproductive assays, we put 10 wild-type DGRP males with 10 *spa* males and 10 *spa* females, counted the number of wt and *spa* offspring after 16 days and calculated the relative number of wt offspring. We measured reproductive success early and late in life (day 4 and day 25). For lifespan, we checked survival 5-6 days a week until all flies died. Finally, for each isoline, we randomly selected 15-20 individuals, dissected the guts on days 5 and 26, and pooled them in groups of five, resulting in 3-4 replicates per isoline for each age group. This figure was created with BioRender.com.

**Figure 2:**
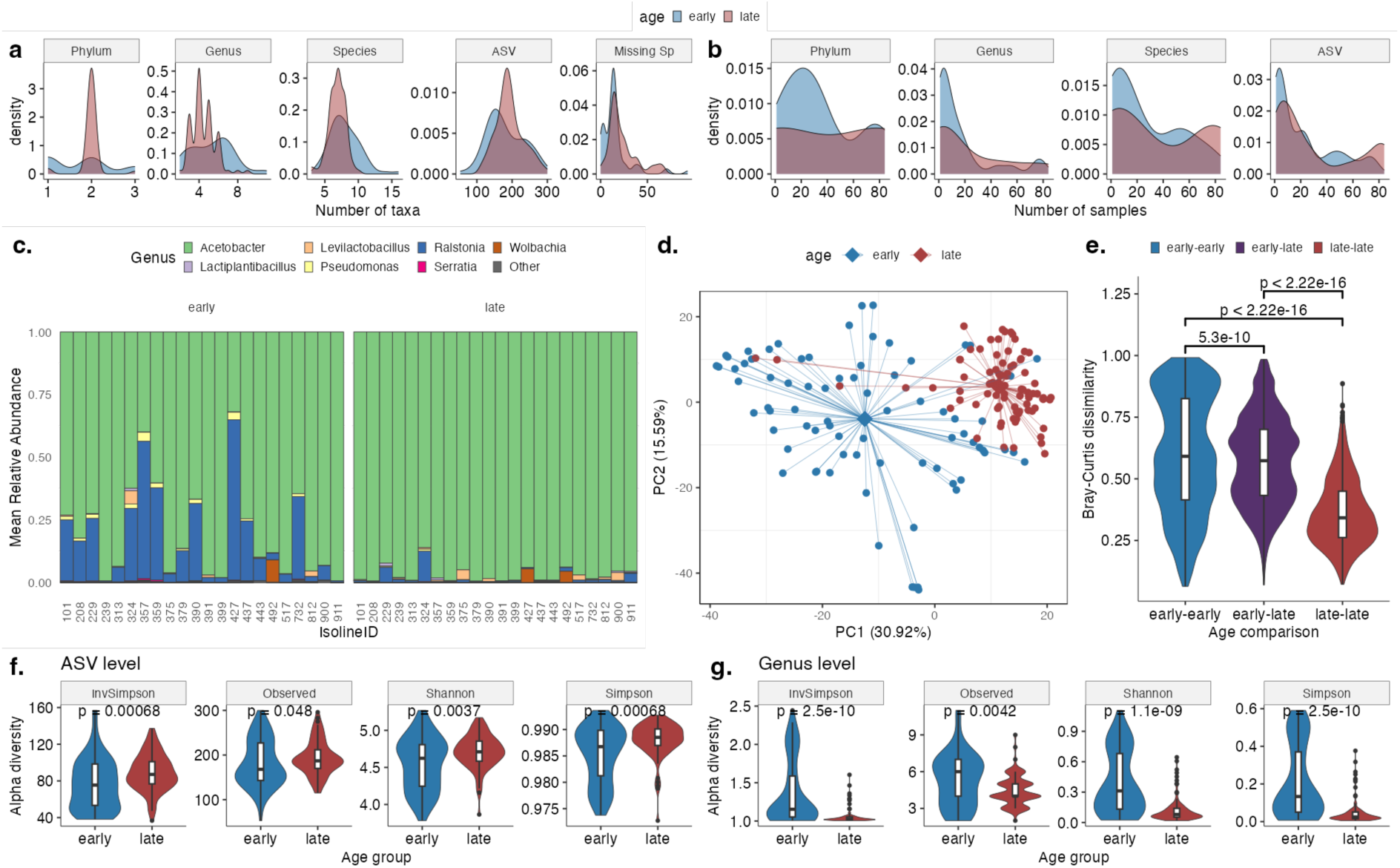
Early and Late Life Microbiota across DGRP Lines. a) Density plots displaying the diversity of taxa across early and late-life samples for various taxonomic ranks. “Missing Sp” indicates the number of ASVs lacking species-level annotations. b) Density plots illustrating taxa prevalence in young (blue) and old (red) samples across distinct taxonomic levels. c) Mean relative abundance of genera across isolines, categorized by age. d) PCA, employing the Euclidean distance on the CLR-transformed abundance matrix (i.e., Aitchison index). Age group centroids are represented by diamonds, and the distances to all samples within the same group are shown by lines originating from the center. e) Distribution of beta-diversity between samples from different age groups, calculated at the ASV level. f) ASV-level and g) genus-level alpha diversity across age groups, evaluated using various indices. p-values shown on the charts correspond to the Wilcoxon test.

We also estimated the climbing speed of each isoline once a week for 4 weeks (i.e., on days 3, 10, 17, and 24) by measuring the climbing speed of each group of males in their experimental vials. We repeated this procedure three times per vial and took the average. Then, we calculated the average across all replicate groups within each isoline as a measure of the climbing speed of each isoline at each time point and estimated functional senescence as the slope of the age-related decline in climbing speed in four weeks.

### Gut dissection, bacterial DNA isolation and sequencing

For gut dissections, 15-20 flies per isoline were sacrificed right after the two reproduction time points early and late in life. The flies were taken from two vials per isoline, the remaining unsacrificed flies in the vials were transferred to other vials of the isoline to control for density. Dissection of each fly was done separately inside PBS droplets under the microscope by using sterilized forceps. Isolated guts were collected in groups of five to have 3-4 biological replicates per isoline and immediately flash-frozen in liquid nitrogen prior to DNA extraction. DNA extraction from gut tissue was performed with the JetFlex ™ Genomic DNA Purification Kit. The DNA was quantified with Nanodrop-1000 Spectrophotometer (Thermo Scientific, Wilmington, DE) and then sent for sequencing. DNA obtained from gut was used as a template for amplification of the V3-V4 region of bacterial 16S rRNA genes and sequenced using the Illumina MiSeq technology at the Foundation for the Promotion of Sanitary and Biomedical Research (FISABIO), Valencia, Spain.

### Processing of the 16S Amplicon Sequencing Data

We used FastQC [27] to assess the quality and length distribution of the reads supplied by the sequencing center and ensured that the sequences did not contain any adapter or primer sequences. The dada2 pipeline, available as an R package [28], was employed to trim, filter, denoise, merge, and remove chimeras. Based on quality profiles, reads were truncated to 280bp (forward) and 250bp (reverse), followed by filtering. There were two sequencing runs. The first and second runs retained between 94-99.9% and 94-100% of all reads post-filtering.

Errors were estimated independently for the two runs, and subsequently, denoising was performed. Pairs were then merged, resulting in abundance tables. On average, 95% of the filtered pairs from run 1 and 92% from run 2 were successfully merged. The abundance tables from both runs were then combined. We employed the ‘consensus’ method to remove chimeric sequences; that is, samples in a sequence table underwent individual processing for chimeric sequences, with a consensus decision made for each sequence variant. Notably, while 70.4% of unique sequences were chimeric, they accounted for 20.7% of the total reads. For taxonomic assignment, we leveraged the dada2 implementation of the RDP naive Bayesian classifier method described in [29] and the Silva taxonomic database (v138) [30]. This enabled the successful assignment of phylum-level information to 85.4% of all ASVs, and species-level information to 49.35% of all ASVs. Subsequently, we employed the DECIPHER R package [31] to construct a multiple alignment. Using the phangorn R package [32,33], a phylogenetic tree was created based on a Generalized Time Reversible with Gamma rate variation (GTR+G+I) maximum likelihood tree, using a neighbor-joining tree as the starting reference.

#### ASV Filtering

Prior to ASV level filtering, we identified 2020 ASVs. We employed four sequential criteria for the quality filtering of ASVs: **1)** We used only ASVs with lengths ranging from 400 to 465 bp, resulting in 1672 ASVs. **2)** We selected ASVs with phylum-level taxonomic assignment, narrowing it down to 1593 ASVs. **3)** We excluded ASVs that were phylogenetically distant from all other ASVs, leaving 1523 ASVs. **4)** We excluded taxa that are present in less than 1% of all samples, leaving 685 ASVs only. The ASV length distribution, taxonomic assignment rate, and phylogenetic tree both before and after ASV filtering can be viewed in **Supplementary Figure S1**. We further explored whether any DNA extraction or sequencing batches correlated with the presence of ASVs that were filtered out. Some DNA extraction batches contained a higher amount of these ASVs **(Supplementary Figure S2)**. This suggests that these ASVs could indeed be technical artifacts.

### Sample Filtering

We observed significant variations in library sizes across the samples. Four samples with fewer than 10,000 reads were excluded from the analysis **(Supplementary Figure S3)**. We retained only those isolines for which we had at least three replicate samples at both the early- and late-life time points, yielding 161 samples. Given the pronounced disparities in library size, we standardized the number of reads by rarefaction to 10,000 reads for each sample to enable a more equitable comparison of diversity and abundance. While we present the results using only one such rarefied table (unless indicated otherwise), to ensure our results weren’t skewed by a single non-representative rarefaction, we generated 100 distinct rarefied tables and conducted our analysis on each.

### Analysis of 16S Amplicon Sequencing Data

#### Alpha and Beta Diversity

We employed the “estimate_richness” function from the phyloseq package [34] to compute alpha diversity and the distance function for beta diversity calculations. Our analysis incorporated five alpha diversity metrics: “Observed” (number of observed taxa) and three metrics accounting for both richness and evenness: “Shannon”, “Simpson”, and “InvSimpson” (Inverse Simpson). For beta diversity, we used the Bray–Curtis distance. The number of taxa across samples and the prevalence of each taxon (i.e., the number of samples in which each taxon occurs) were determined based on the abundance matrix and non-zero occurrences.

#### PCA and PCoA on microbial abundances

We conducted a PCA on CLR-transformed abundance values using the prcomp function from base R and the transform function within the microbiome R package [35]. Additionally, PCoA plots were generated using the ordinate function from the phyloseq R package [34], based on the Bray-Curtis distance matrix derived from log10(count+1) transformed data.

### Age and life history prediction using microbial abundances

We used “caret” R package [36] to train a random forest model to predict age class (i.e. “early” vs. “late”) or life history traits (i.e. early climbing speed, late climbing speed, functional aging, early reproductive success, reproductive aging, F1 quality, lifespan) using species level CLR transformed abundance data (number of features = 27). We used 5 times repeated 5-fold cross-validation to train the model. Importantly, train and test data included a non-overlapping set of isolines, preventing data leakage. We trained the model on 15 isolines and their 111 samples and tested it on 7 isolines, and their 50 samples. Accuracy was calculated using “confusionMatrix” function for age prediction. MAE and Rsq values were used for the performance evaluation of regression models with life history traits. We used the “varImp” function from the “caret” package to identify the most important features (i.e. species).

Because our primary goal was to evaluate whether microbiota can predict age or life history traits, not to deploy a predictive model, we sought to ensure that the small sample size and random train/test splits did not undermine our conclusions. Consequently, we repeated the entire procedure 20 times with new train–test splits, resulting in 20 different models. We reported the performance range of all models in the results section and all results are given as **Supplementary Tables 1 and 2**.

### Age and isoline effects on gut microbiota composition

We used DESeq2 R package [37] to analyze differential ASV abundances between early- and late-life and between isolines. We used the ASV abundances rarefied to 10,000 reads per sample and analyzed only the ASVs that are found in at least 5 samples in early- and late-life and have on average 1 in 10,000 abundance across early- and late-life samples. We used a model that takes sequencing run, DNA batch, age, and isoline into account and we tested for the significance of age and isoline separately, using a likelihood ratio test. ASVs with FDR adjusted p value < 0.1 are considered significant.

### Linking life history traits to gut microbiota composition

We collected all ASVs that show significant differences between isolines, aggregated ASV level information to species level information, and then tested for association between their abundances and life-history traits, using the Spearman correlation test. We calculated the mean abundance level of each bacterial species for each isoline and age category and used these values to correlate with life history. P-values are corrected using the FDR procedure as implemented in “p.adjust” function in base R and an FDR-adjusted p-value < 0.1 is considered as a significant association.

## Results

Using *Drosophila* Genetic Reference Panel (DGRP) isolines, we explored the association between various life history traits and gut microbiota. Following the quality filtering of microbiota samples (see Methods), we analyzed 22 isolines. We investigated five key traits related to life-history and physiological performance: average lifespan, early climbing speed, functional aging (defined as the decline in climbing speed with age), early reproductive success, and reproductive senescence (measured as the difference between late and early reproductive successes). The experimental design is outlined in **Figure 1** and life history traits associated with these 22 isolines are provided as **Supplementary Table 3**.

We noted considerable variation in life history traits across isolines. For instance, the longest recorded lifespan was 2.3 times the shortest lifespan. Similarly, the highest early climbing speed was 3.6 times the lowest value, and the highest recorded early reproductive success was 6.16 times the lowest. The mean lifespan stood at 37.8 days with a standard deviation of 8.23 days. A comprehensive set of summary statistics can be found in **Supplementary Table 4**.

We further analyzed the correlations between the life history traits. Except for early climbing speed and functional aging **(Supplementary Figure S4h)**, we did not observe significant correlations **(Supplementary Figure S4)**.

### Diversity of Gut Microbiota across DGRP Lines in Early and Late Life

After quality filtering for Amplicon Sequence Variants (ASV) and samples and performing read rarefaction (see Methods), we identified 677 ASVs across 161 samples. Initially, we analyzed the distribution of detected taxa across samples, at different taxonomic levels **(Figure 2a)**.

Samples had a median of 184 ASVs, ranging between 53 and 300. From this pool, on average 20 ASVs (median: 14, range: 0-80) remained unidentified at the species level. The median counts for recognized species, genera, and phyla were 7 (range: 3-16), 5 (range: 2-11), and 2 (range: 1-3), respectively. These findings align with the literature emphasizing that *Drosophila* gut microbiota mainly comprises a select array of bacterial species, typically between 5 to 20 [38]. We also observed a prominent variation across isolines, with the most diverse isoline hosting twice as many taxa as the least diverse one at nearly all taxonomic tiers **(Supplementary Table 5)**.

Further, we examined the prevalence of taxa at each taxonomic level. ASVs were present in an average of 44.5 out of 161 samples, species in 48, genera in 30.4, and phyla in 75.5 samples **(Figure 2b)**. Notably, even at this general level, we observed differences between young and old samples. Older samples exhibited slightly reduced species and genera diversity, whereas ASV diversity increased **(Figure 2a)**. Distribution of taxa prevalence also varied between the young and old groups, with an increased number of species and ASVs with higher prevalence in older samples **(Figure 2b)**. We further analyzed the taxa prevalence across isolines and found that while many taxa were specific to certain isolines (shared in 1 to 5 isolines), some were consistently found across a large number of isolines, especially at the species and ASV levels **(Supplementary Figure S5)**.

Next, we explored the genera composition across isolines and age groups **(Figure 2c)**, noting a decline in diversity and relative abundance of certain taxa, such as *Ralstonia*, with age. Interestingly, *Wolbachia* was only detected in two isolines: one (isoline 427) only in old flies, and one (isoline 492) both in young and old groups. PCA on CLR transformed data highlighted age-centric clustering (predominantly in the first principal component), which explained 30.92% of the data variance **(Figure 2d)**. Similarly, PCoA conducted on log-transformed abundance data showed age-based clustering, independent of sequencing batches **(Supplementary Figure S6)**.

We then analyzed alpha diversity differences between young and old samples at both ASV and genera levels **(Figure 2f-g)**. Our analysis validated the observed contrasting diversity trends between genera (decreasing with age) and ASV (increasing with age) levels in **Figure 2a**. It’s pertinent to mention that our samples were pooled, not derived from individual flies, signifying the alpha diversity here represents community-level diversity. Lastly, we examined the beta diversity between young and old samples, utilizing the Bray-Curtis dissimilarity measure **(Figure 2e)**. A marked difference emerged between young and old samples, with older samples showing more similarity amongst themselves than the younger samples.

### Prediction Models

Given the large variation in gut microbiota and life histories, next, we asked whether we could predict age or life history traits using gut microbiota. We first trained a random forest model on 15 isolines to predict age (early vs. late) using CLR-transformed species level abundance information. We tested the model on the remaining non-overlapping 7 isolines and achieved 84% accuracy, whereas the no information rate, which reflects the accuracy one would get by always predicting the most frequent class, was 54% **(Figure 3a)**. The most important species contributing to the classification were *Acetobacter tropicalis*, *Ralstonia pickettii*, *Acetobacter indonesiensis*, *Leuconostoc pseudomesenteroides*, and *Acetobacter persici* **(Figure 3b)**. Then, we trained models to predict life history traits using microbial abundances. While in some train-test split sets, we achieved reasonable performance, in the majority of cases the model failed to predict any of the life history traits (R square, mean absolute error, and no information rate mean absolute errors are given as **Supplementary Table 2**). To ensure our results with age prediction are robust, we trained 20 models for age prediction with differing train - test splits. We achieved an average (median) accuracy of 86.27% with a minimum of 72.92 and a maximum of 96% **(Supplementary Table 1**). Moreover, *Acetobacter tropicalis* and *Ralstonia pickettii* were consistently the two most important features across all models, while the remaining three were on average half of the time among the top 5 most important species. We conclude that gut microbiota can predict the age of samples with high accuracy, but not the life history traits of the isolines, at least with this sample size.

**Figure 3:**
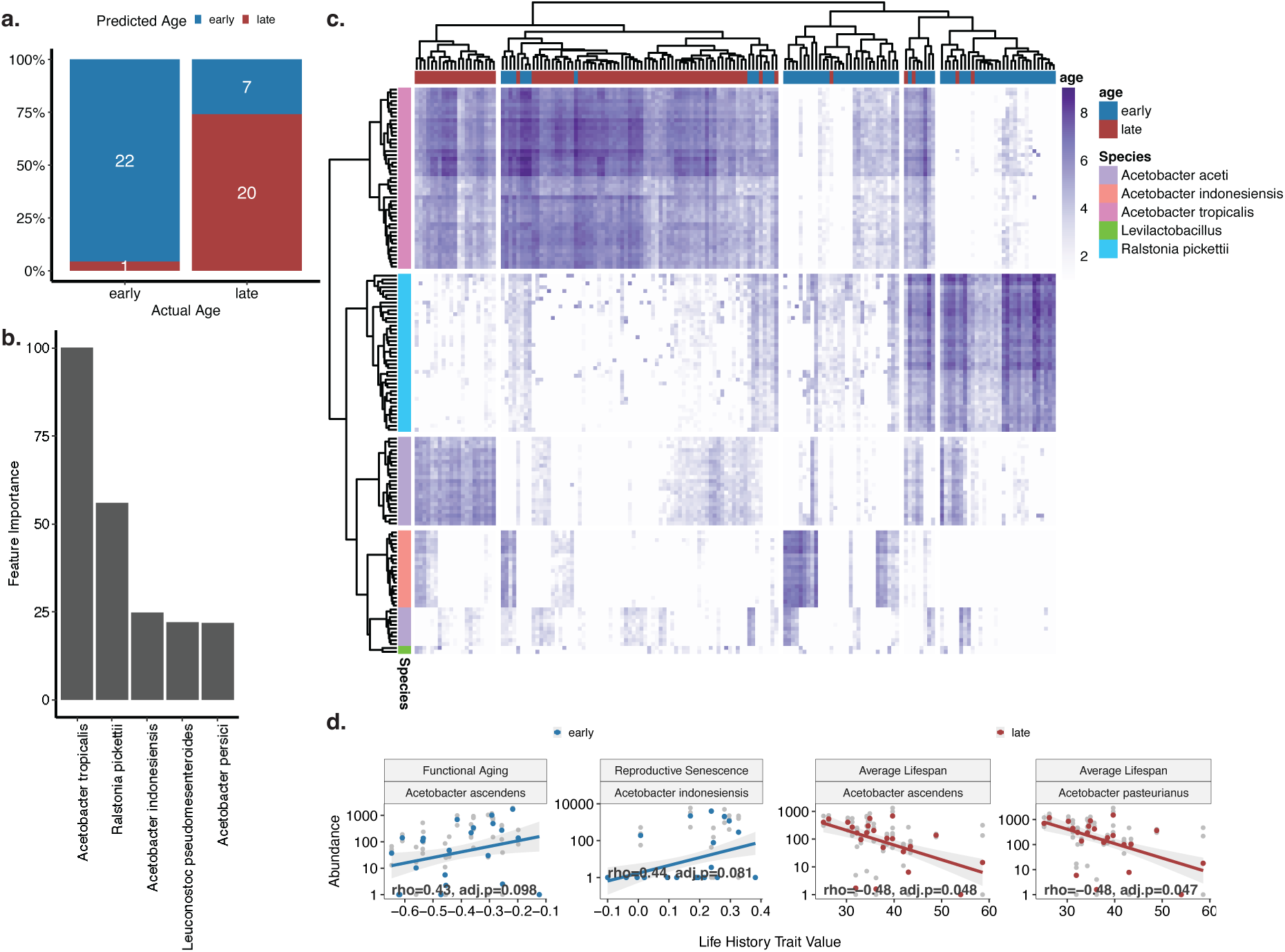
Male age- and life-history-associated taxa in *Drosophila* gut. a) Performance of age-prediction using CLR transformed species abundances and a random forest model. b) Most important features in random forest model for prediction of age. c) ASVs showing differential abundance with age and their species annotations. d) Significant correlations (FDR-adjusted p-value<0.1) between male life history traits and abundance values of species that show significant differences between isolines (FDR-adjusted p-value<0.1). Gray points indicate individual samples; colored points represent the mean value for each isoline.

### Age-related Changes in Gut Microbiota and Life History Traits

We found a significant effect of male age, isoline, and their interaction on bacterial abundances. Although there was an overall increase in ASV diversity with male age, the direction of age-specific change in bacterial diversity differed with respect to different isolines. In general, the relative abundance of *Acetobacter* genus increased with age, while other genera such as *Ralstonia* became less abundant **(Figure 3c)**.

When we explored the link between life history traits and bacterial diversity, we observed some interesting trends. In particular, some trends were in opposite directions between early and late samples, however, none of the associations were significant after correcting for multiple testing **(Supplementary Figure S7)**.

We observed significant associations (FDR adjusted p-value < 0.1) between abundances of different *Acetobacter* species and life history traits. In early life, *A. ascendens* abundance was positively correlated with functional aging and *A. indonesiensis* abundance was positively correlated with reproductive senescence. Finally, late-life abundances of *A. ascendens* and *A. pasteurianus* were both negatively correlated with average lifespan **(Figure 3d)**.

### Testing the potential influence of rarefaction on results

Since we observed a large variation across library sizes, we rarefied all samples to 10,000 reads before all analyses. Since random sampling of 10,000 reads may bias the results, we then performed rarefaction 100 times and made all calculations. We first checked the alpha diversity differences between early- and late-life samples and confirmed that the trend is observed across all rarefactions **(Supplementary Figure S8)**. Then we checked differential abundances, following the exact same procedure described above, repeated for 100 independent rarefactions. The 5 species reported in the main text that have differential abundance between early- and late-samples were detected as differentially abundant in all 100 rarefactions **(Supplementary Table 6)**. There were two more species *Pseudomonas yamanorum* and *A. persici* that were significant in 30 and 9 of the rarefactions, respectively, but with only 1 or 2 ASVs. We repeated the same analysis for the associations between species and life history traits **(Supplementary Table 7).** Three of the four associations reported in **Figure 3d** were also detected in the same direction in all 100 rarefactions, while the association between *A. ascendens* in early life and functional aging was observed only in 57 of the associations. A negative association between *P. yamanorum* and reproductive senescence was also observed in both early- and late samples but only in a limited number of rarefactions, 7 and 1 out of 100 rarefactions, respectively.

## Discussion

In this study, we explored the link between gut microbiota and male life history traits. First, as previously shown in the literature, we found age-related changes in male gut microbiota diversity and composition, with species and genera diversity decreasing with age and *Acetobacter* becoming dominant in the guts of older individuals, in line with the literature in males [23]. Second, we identified significant associations between gut microbiota composition and several male life history traits. In particular, the abundance of different species of *Acetobacter* was negatively correlated with lifespan and positively correlated with reproductive and functional aging, suggesting a potential negative impact of these species on male aging and reproduction in fruit flies. Third, we showed that gut microbiota composition could robustly predict the chronological age of male flies while failing to predict their life history traits. This suggests that there are consistent age-related shifts across genotypes, while microbiota-life history trait relationship is more complex, requiring individual-level information or higher sample size.

### Age-Related Changes in Male Gut Microbiota

We found that ASV-level bacterial diversity was higher, while species and genus diversity was lower, in older male flies. In humans, studies present mixed findings on how alpha diversity changes with age. One study reported a decrease in alpha diversity with age [39], while another observed an initial increase followed by a decline at very old ages [40]. Conversely, some found that bacterial diversity in the gut increased with age [41], while others found no significant effects of age on alpha diversity [42,43]. The differences in genetic background, culture, and lifestyle are suggested as potential reasons for the inconsistencies in human studies [39]. In male fruit flies, previous findings showed that as flies age OTU diversity is higher [11] while species diversity is lower [23] aligning with our results.

The observed increase in ASV-level richness alongside a decline in species-level richness may reflect a rise in unidentified ASVs at an old age. However, it is difficult to determine whether this pattern results from limited database annotations, possible within-host evolution (despite the short 21-day timespan), or other technical factors. In general, the difference between human and fly studies can be because flies are generally kept under controlled conditions during aging while this is not possible in humans. As a result, it is harder to dissociate age effects from other factors such as diet in human studies compared to flies.

We found that the relative abundance of *Acetobacter* was higher in older individuals, in line with previous findings [23,44]. This was accompanied by a decline in the abundance of other species, such as those from the genus *Ralstonia,* with age. Although *Ralstonia* has been identified previously in the fruit fly gut in relatively small amounts [44], its higher abundance in young individuals in our study suggests it may play an important role in some fly strains, including the DGRP lines.

It is important to mention that we pooled 5 genetically identical flies (DGRP isolines), allowing us to approximate alpha diversity, which refers to the microbial diversity within a single sample or individual [45]. In ecological terms, alpha diversity reflects how many different taxa coexist locally and how evenly they are distributed, offering insights into community complexity and resilience. Higher alpha diversity is often associated with ecological stability and functional redundancy, which may buffer the host against environmental or physiological stressors—an especially relevant consideration in the context of aging, where declining resilience is a common theme [46]. Considering that these individuals are genetically identical and were reared together, their microbiota can be considered as a single community. Still, stochastic events might have caused genetically identical individuals to have slightly different gut microbiota composition. Therefore, our diversity measure can also be considered as gamma diversity which is the total diversity observed across multiple individuals or habitats [47]. Gamma diversity in this context provides a snapshot of the overall microbial repertoire within a genetically homogeneous group, and how it may change over time with host aging.

We also examined beta diversity, which captures variation in microbial community composition between samples. Interestingly, older flies had more similar gut microbiota to each other compared to young flies. In ecological terms, beta diversity reflects the extent of differentiation between communities and is often shaped by dispersal, selection, and drift. A decline in beta diversity may suggest a convergence toward a limited set of taxa, possibly reflecting ecological filtering or reduced niche space in the aging gut [48]. This pattern contrasts with findings from human studies, where beta diversity typically increases with age [49,50]. One possible explanation for this unexpected pattern is technical. Since our samples were pooled from multiple individuals, the reduced variability may reflect a regression toward the mean. Biologically, the lower inter-individual variability in older flies might also result from their rearing under highly controlled conditions with identical diets throughout aging - conditions that differ markedly from the more variable environments experienced by aging humans [39], or wild flies in nature. Alternatively, this may reflect a convergence toward a simplified community dominated by a few taxa, such as *Acetobacter*, in late life. This microbial streamlining could represent a loss of ecological complexity and reduced functional flexibility, potentially contributing to age-associated physiological decline.

### Linking Gut Microbiota with Male Life History Traits

We found no correlation between bacterial diversity and male life history traits. Previous studies in humans have generally linked greater microbial diversity to better health [51]. However, the literature has mixed findings regarding the impact of bacterial diversity on male life history in flies, with most research focusing on lifespan rather than reproduction. While a more diverse microbiome has been linked to increased lifespan in both sexes [52], experiments with axenic flies showed that introducing multiple bacterial species progressively reduced lifespan, with axenic flies living the longest and those inoculated with five species having the shortest lifespans [17]. The absence of a link between bacterial diversity and male life history traits in our study could be due to our limited statistical power, given the relatively small sample size (i.e., 22 isolines). Alternatively, it is possible that bacterial diversity does not significantly influence male life history in fruit flies. In fact, a recent study found that feeding old male fruit flies with the gut microbiome of young flies affects neither locomotor activity nor lifespan [11], suggesting the limited ability of gut microbiome to modulate male life history in this species. Further studies with larger sample sizes are needed to explore these possibilities.

Although bacterial diversity was not associated with male life history, we did find specific links between the abundance of certain bacterial species and aging-related traits. In particular, certain species of *Acetobacter* (i.e., *A. ascendens*, *A. indonesiensis* and *A. pasteurianus)* were linked to faster actuarial, functional or reproductive aging. These findings contrast with Wesseltoft et al (2024) that showed transferring young microbiota into old males had no effect on lifespan or locomotor activity. Several factors can explain this difference. First, despite the presence of beneficial bacterial species in young guts, the aged guts of old flies may already be dysfunctional, rejecting the young microbiota and limiting its effect on life history (i.e., host-mediated effects, see [53]). Second, transplantation of beneficial microbiota might be more effective earlier in life, before aging-related damage has already accumulated [10]. Finally, it is noteworthy that the observed correlations between certain bacterial species and life history traits in our study indicate associations rather than causation. Additional research is needed to determine the functional roles of these species, their interactions with the host, and the underlying mechanisms through which they may influence lifespan.

Our findings linking late-life *A. indonesiensis* abundance to reproductive senescence highlight the potential negative effects of this species on male reproduction. A previous study demonstrated that males mono-associated with *A. pomorum* had shorter mating duration and lower number of offspring compared to those mono-associated with *L. plantarum* [20]. While *Acetobacter* species are generally associated with benefits to female reproduction [17,54,55], our findings, together with those of Morimoto et al. (2017), suggest that certain species of *Acetobacter* may have an adverse effect on male reproduction. Further research is required to investigate additional bacterial species and to examine male reproductive success in more competitive environments.

Finally, we also found correlations between the abundance of *A. ascendens* and *A. pasteurianus* with ageing, specifically faster functional ageing and shorter lifespan. A previous correlational study by Walters et al. (2020) also linked *Acetobacter* abundance to faster ageing [56]. They observed that female flies from low-latitude populations had shorter lifespans, higher early reproduction, and more acetic acid bacteria (including *Acetobacter*), while the ones from high-latitude populations exhibited longer lifespans, lower early reproduction, and fewer acetic acid bacteria. Likewise, Gould et al (2018) found that bacterial combinations that caused short lifespan in both sexes led to high fecundity, and combinations that caused long lifespan resulted in a low fecundity. These findings suggest that gut microbiota composition can drive a life history trade-off between lifespan and reproduction in females. However, since male reproductive success was not investigated in these studies, it remains unclear whether a similar trade-off exists in males. Given that we found no positive effects of *Acetobacter* species in males, it is also possible that certain gut microbiota compositions promote high fecundity and short lifespan in females while conferring no benefit to males, potentially due to sexually antagonistic selection [57].

## Conclusion

Our study reveals that gut microbiota composition in male *D. melanogaster* undergoes significant age-related changes, with *Acetobacter* species becoming increasingly dominant in late life and correlating with shorter lifespan, faster functional decline, and reproductive aging. While microbial diversity patterns shifted across taxonomic levels with age, we found no consistent associations between overall diversity and male life history traits. Notably, gut microbiota composition accurately predicted chronological age but failed to predict life history traits, suggesting that while age-associated microbial shifts are robust, the current experimental setup lacks the resolution or power to capture microbiota-life history links. These findings underscore the need for future mechanistic and longitudinal studies to uncover the causal roles of specific microbial taxa in shaping male aging and reproductive strategies.

## Data and Code Availability

Raw sequencing data have been deposited in the European Nucleotide Archive (ENA) at EMBL-EBI under accession number PRJEB88786. Life history trait data is available in **Supplementary Table 3.** All processed data, including the data underlying each figure is available in BioStudies with **accession number S-BSST2029**. All code for preprocessing, statistical analysis, and visualization is available at https://github.com/mdonertas/DGRP_16S_MaleLH.

## Acknowledgment

This study was supported by the Spanish Ministry of Economy and Competitivity grants: PGC2018-099344-B-I00 (AL), CGL2014-58722-P (PC and JILL), RYC-2013-12998 (PC), and RYC-2012-11872 (JILL). HMD is funded by Carl-Zeiss-Stiftung (P2021-00-007). PA was supported by FCT through a PhD grant (number 2021.05611). ZS was supported by the Atracció de Talent Fellowship and the Leverhulme Trust Early Career Fellowship (ECF-2022-214).

