## Supplemental Figures for "Age-dependent gut microbiota dynamics and their association with male fitness traits in *Drosophila melanogaster*"

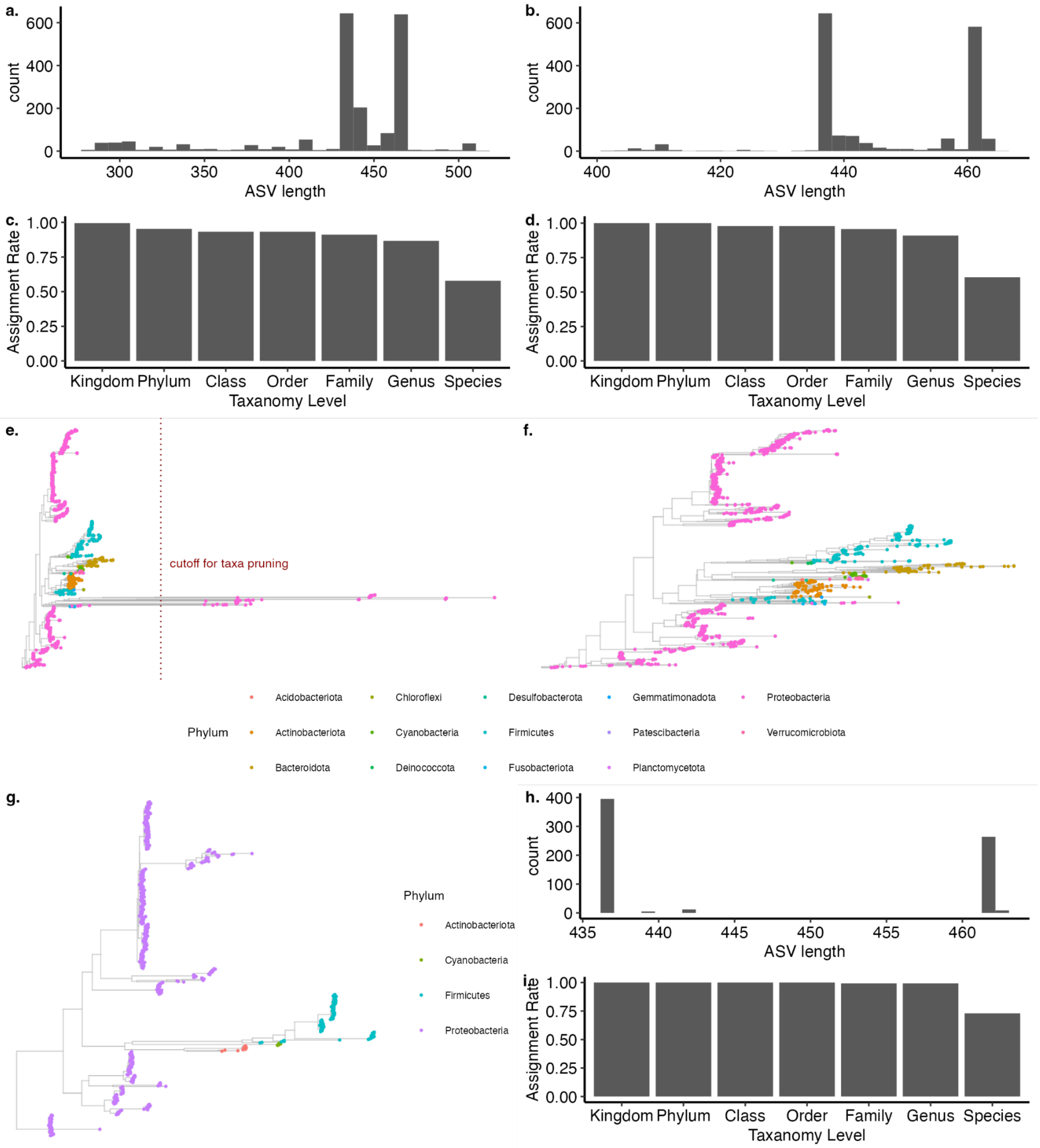


**Supplementary Figure S1:** a) ASV length distribution before any filtering. b) ASV length distribution after the first filter, i.e. selecting ASVs within 440 to 465 bp length. c) Taxa assignment rate after the first filter. d) Taxa assignment rate after the second filter, i.e. excluding the ASVs without phyla assignment. e) Phylogenetic tree after the second filter. f) Phylogenetic tree after the third filter, i.e. excluding divergent taxa. g) Phylogenetic tree after the forth filter, i.e. excluding ASVs that are only present in one sample. h) ASV length distribution after the forth filter. i) Taxa assignment rate after the third filter.

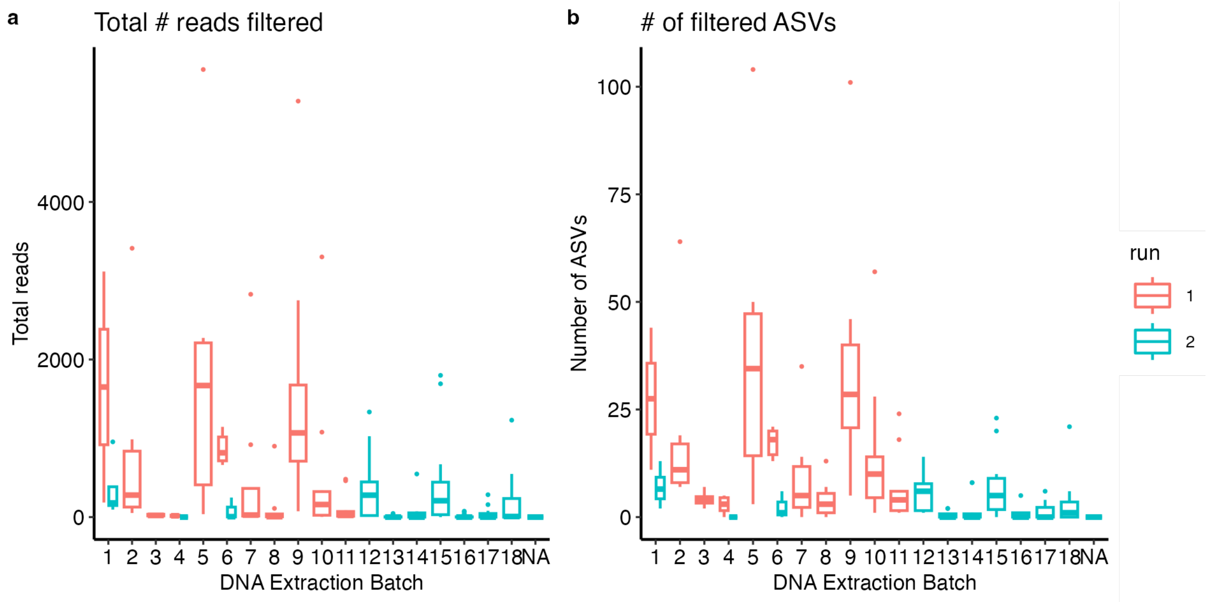


**Supplementary Figure S2:** The total number of reads (a) and unique ASVs (b) filtered out based on ASV exclusion criteria (i.e. ASV length, phyla assignment, and phylogenetic divergence).

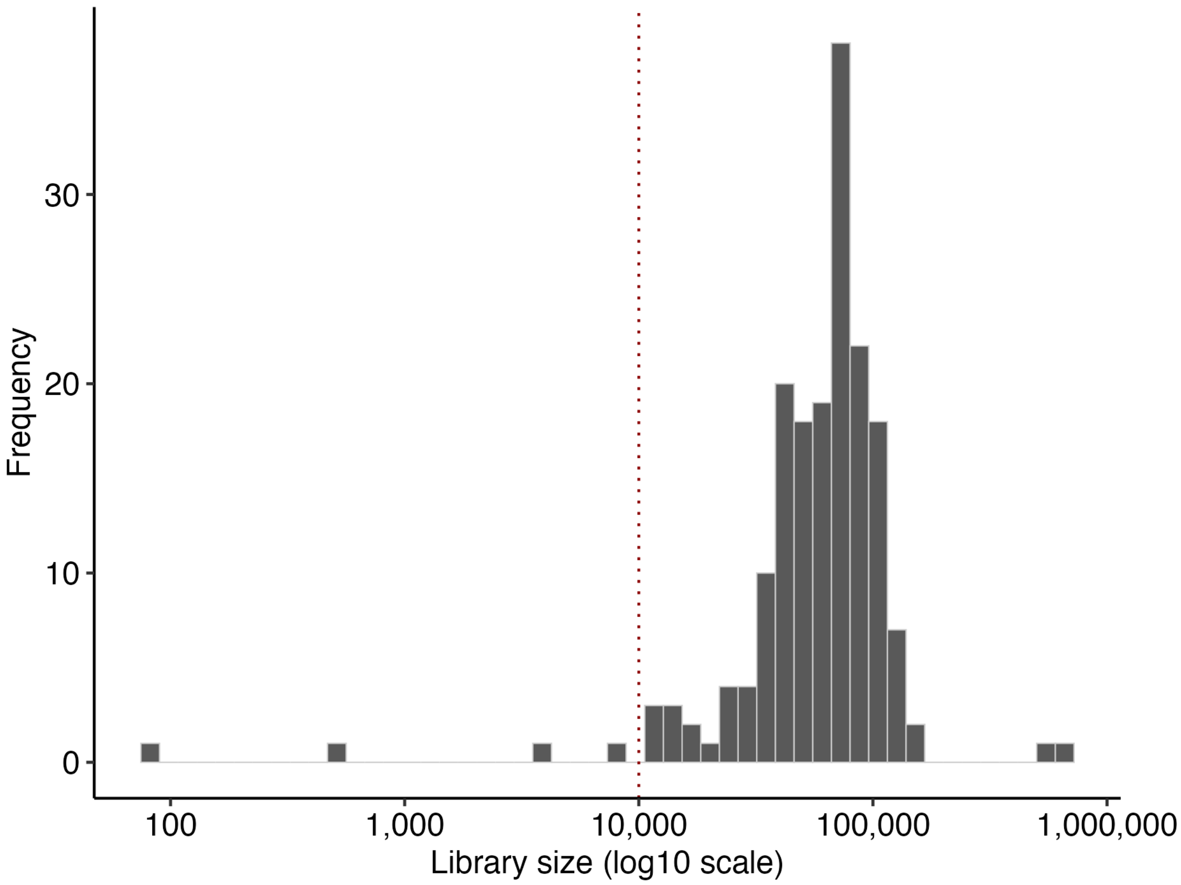


**Supplementary Figure S3:** Distribution of library size (x-axis, log 10 scale) across samples.

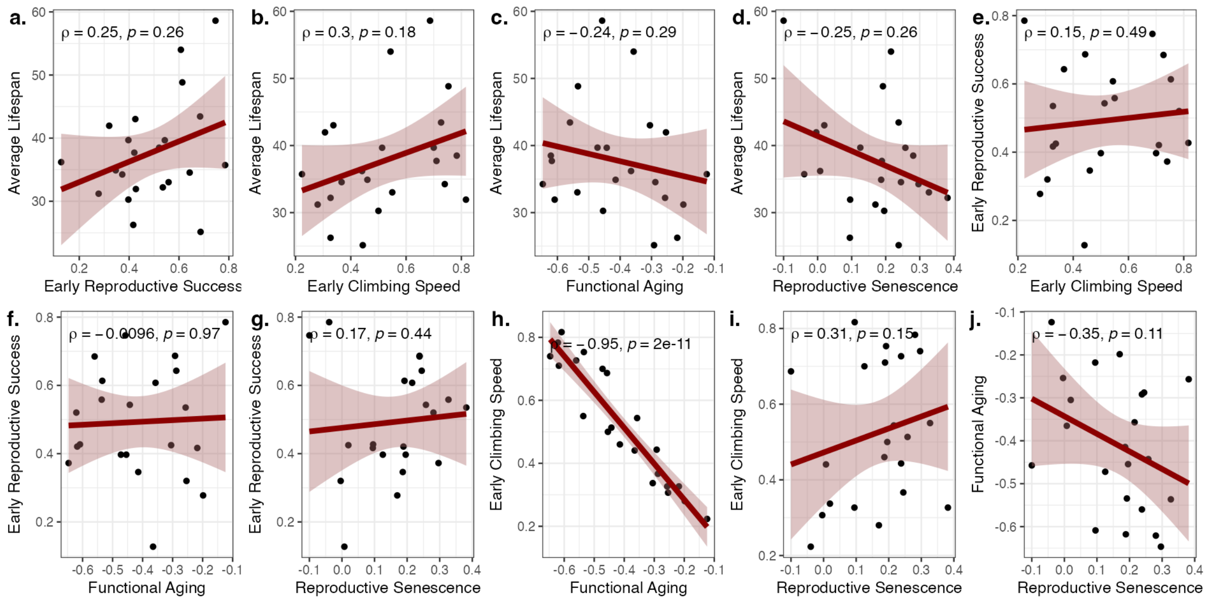


**Supplementary Figure S4:** Distributions and correlations across life history traits.

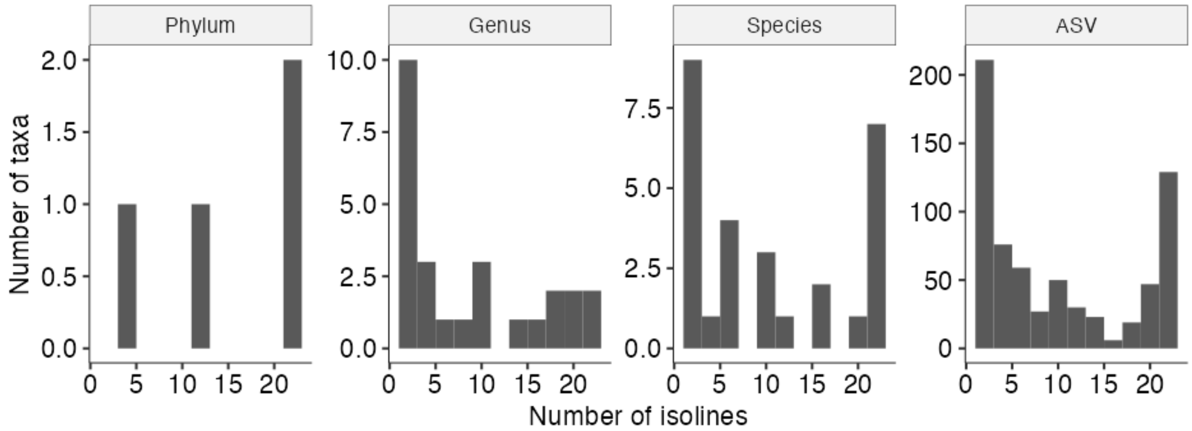

**Supplementary Figure S5:** Isoline prevalence of taxa across different taxonomic levels.

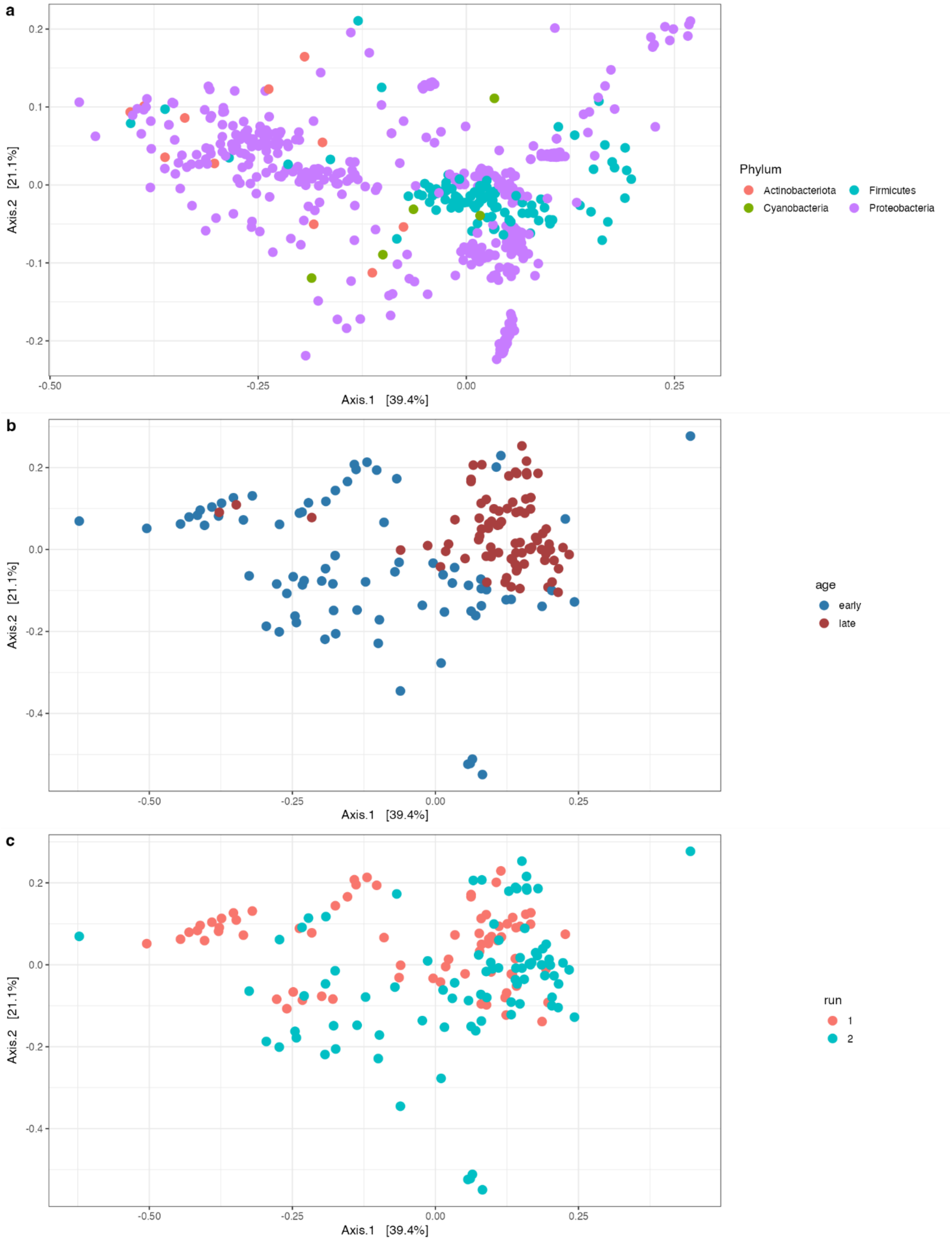


**Supplementary Figure S6:** PCoA calculated using Bray-Curtis distances between samples based on log10(count + 1) transformed abundance data, highlighting a) ASV coordinates colored by phyla, b) samples colored by age c) samples colored by sequencing run.

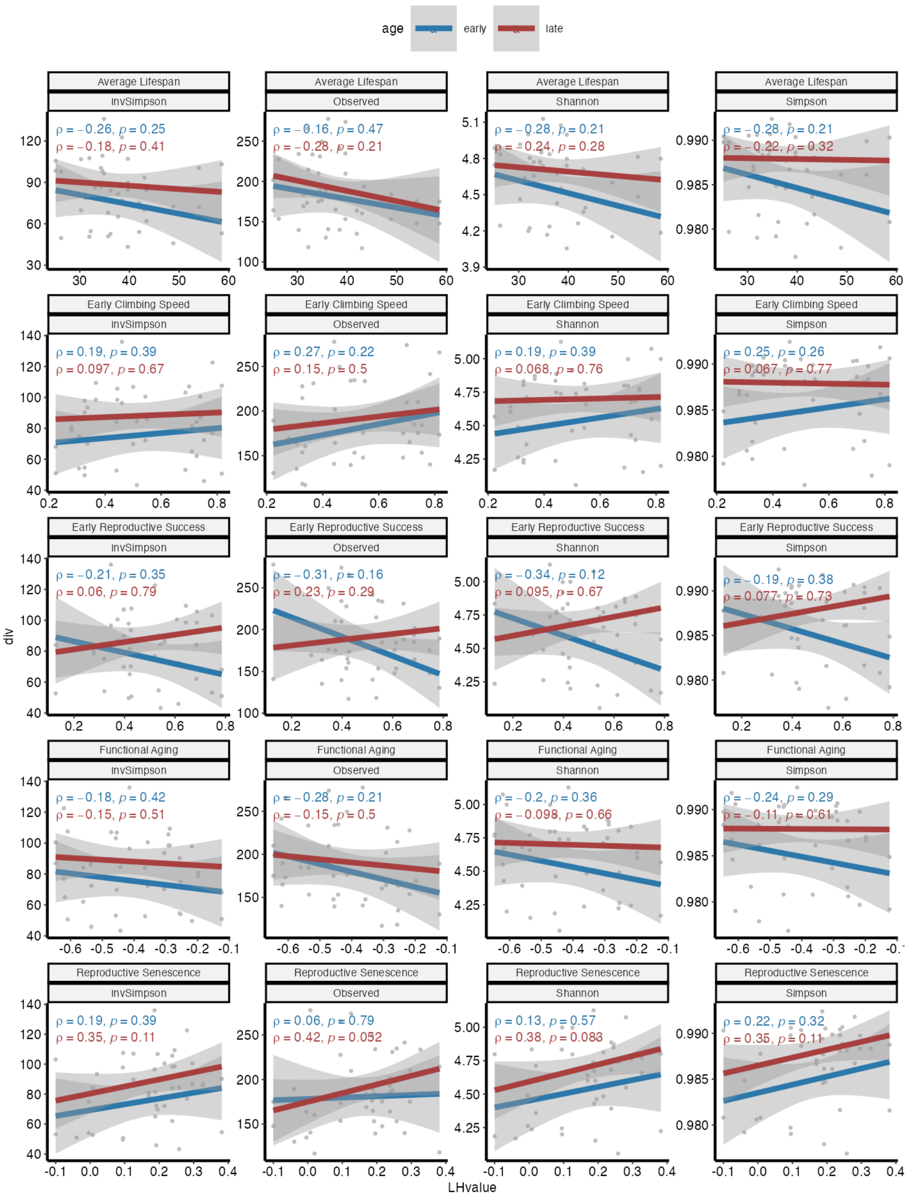


**Supplementary Figure S7:** Associations between alpha diversity measures in early- and late-samples and life history traits.

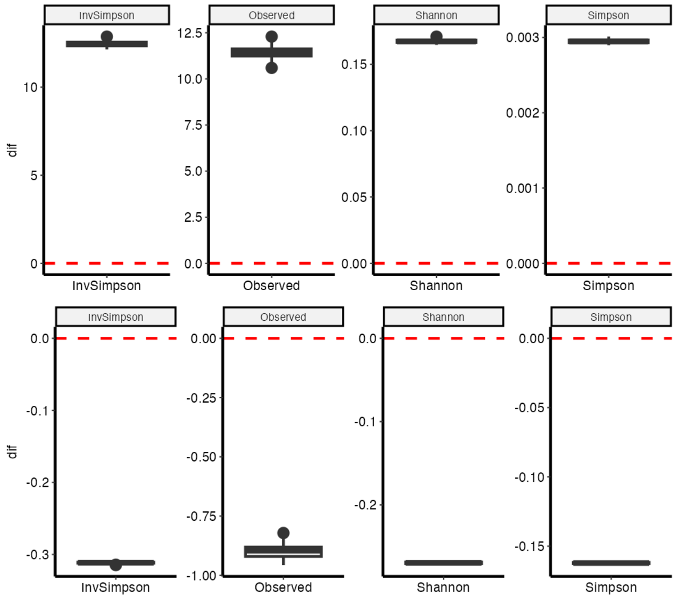


**Supplementary Figure S8:** Difference between alpha diversities of young and old samples across 100 rarefactions to 10,000 reads.
